# 3D Microwell Platforms for Control of Single Cell 3D Geometry and Intracellular Organization

**DOI:** 10.1101/2020.07.19.209460

**Authors:** Robin E. Wilson, Aleksandra K. Denisin, Alexander R. Dunn, Beth L. Pruitt

## Abstract

**Introduction:** Cell structure and migration is impacted by the mechanical properties and geometry of the cell adhesive environment. Most studies to date investigating the effects of 3D environments on cells have not controlled geometry at the single-cell level, making it difficult to understand the influence of 3D environmental cues on single cells. Here, we developed microwell platforms to investigate the effects of 2D vs 3D geometries on single-cell F-actin and nuclear organization.

**Methods:** We used microfabrication techniques to fabricate three polyacrylamide platforms: 3D microwells with a 3D adhesive environment (*3D/3D*), 3D microwells with 2D adhesive areas at the bottom only (*3D/2D*), and flat 2D gels with 2D patterned adhesive areas (*2D/2D*). We measured geometric swelling and Young’s modulus of the platforms. We then cultured C2C12 myoblasts on each platform and evaluated the effects of the engineered microenvironments on F-actin structure and nuclear shape.

**Results:** We tuned the mechanical characteristics of the microfabricated platforms by manipulating the gel formulation. Crosslinker ratio strongly influenced geometric swelling whereas total polymer content primarily affected Young’s modulus. When comparing cells in these platforms, we found significant effects on F-actin and nuclear structures. Our analysis showed that a 3D/3D environment was necessary to increase actin and nuclear height. A 3D/2D environment was sufficient to increase actin alignment and nuclear aspect ratio compared to a 2D/2D environment.

**Conclusions:** Using our novel polyacrylamide platforms, we were able to decouple the effects of 3D confinement and adhesive environment, finding that both influenced actin and nuclear structure.

## Introduction

Cells sense and respond to their mechanical environment through a variety of signaling pathways in a process broadly termed mechanotransduction.^17,26^ Substrate stiffness,^16,57^ patterned spread area and shape,^44,52^ and topography^35^ can all influence cell structure and function. Most of the studies to date investigating the effects of environmental cues on cell structure or function have been done on 2D substrates. These substrates are simple to fabricate, commercially available, and compatible with microscopy, making them convenient for *in vitro* experiments. Such 2D studies have demonstrated controlling a cell’s 2D geometry via protein patterning impacts cytoskeletal structure,^52,55^ contractile force generation,^44^ and differentiation.^40^ However, most cells in the body exist in complex, 3D tissues, indicating that 2D environments cannot fully recapitulate the confinement and mechanics of a tissue. Therefore, we sought to test how dictating a 3D geometry for single cells influences cell morphology and structure.

Recently, there has been growing interest in using 3D microtissues and organoids for disease modeling,^6,15,29^ drug testing,^1,32^ and regenerative medicine.^5,58^ Many of these studies have shown that 3D microtissues can exhibit more physiologically relevant responses to stimuli when compared to cells cultured on 2D substrates, most commonly tissue culture plastic.^37,45,49,54^ However, cells in such 2D and 3D environments experience numerous differences in environmental cues including substrate stiffnesses, adhesion ligand type and accessibility, cell-cell interactions, and morphology. These differences make it difficult to determine whether the geometric confinement and mechanical environment alone can cause changes in cell structure and behavior.^2^

A few recent studies have implemented engineering approaches to construct single-cell microwells used to investigate the effects of more controlled 3D microenvironments on individual cells.^4,38,50^ These studies determined that the shape and geometry of the 3D microenvironment can influence intracellular organization, but they lacked 2D comparisons^4^ and did not utilize fully enclosed 3D microwells.^38,50^ To gain insight into how specific factors like geometry and cell adhesion to the ECM affect intracellular structure, we compared morphologies of single cells in a) 3D microwells with a 3D adhesive environment (*3D/3D*), b) cells adhering to flat, 2D gels with 2D patterned protein as adhesive “islands” (*2D/2D*), and cells in 3D microwells with 2D adhesive “islands” at the bottom only (*3D/2D*). To do so, we created micropatterned polyacrylamide substrates and manipulated both the geometry and protein adhesion areas to create three platform variations above (Figure 1). These platforms enabled unique experiments that could specifically investigate the effects of the 3D geometry and of the 3D cell-ECM adhesions on intracellular structure.

**Figure 1:**
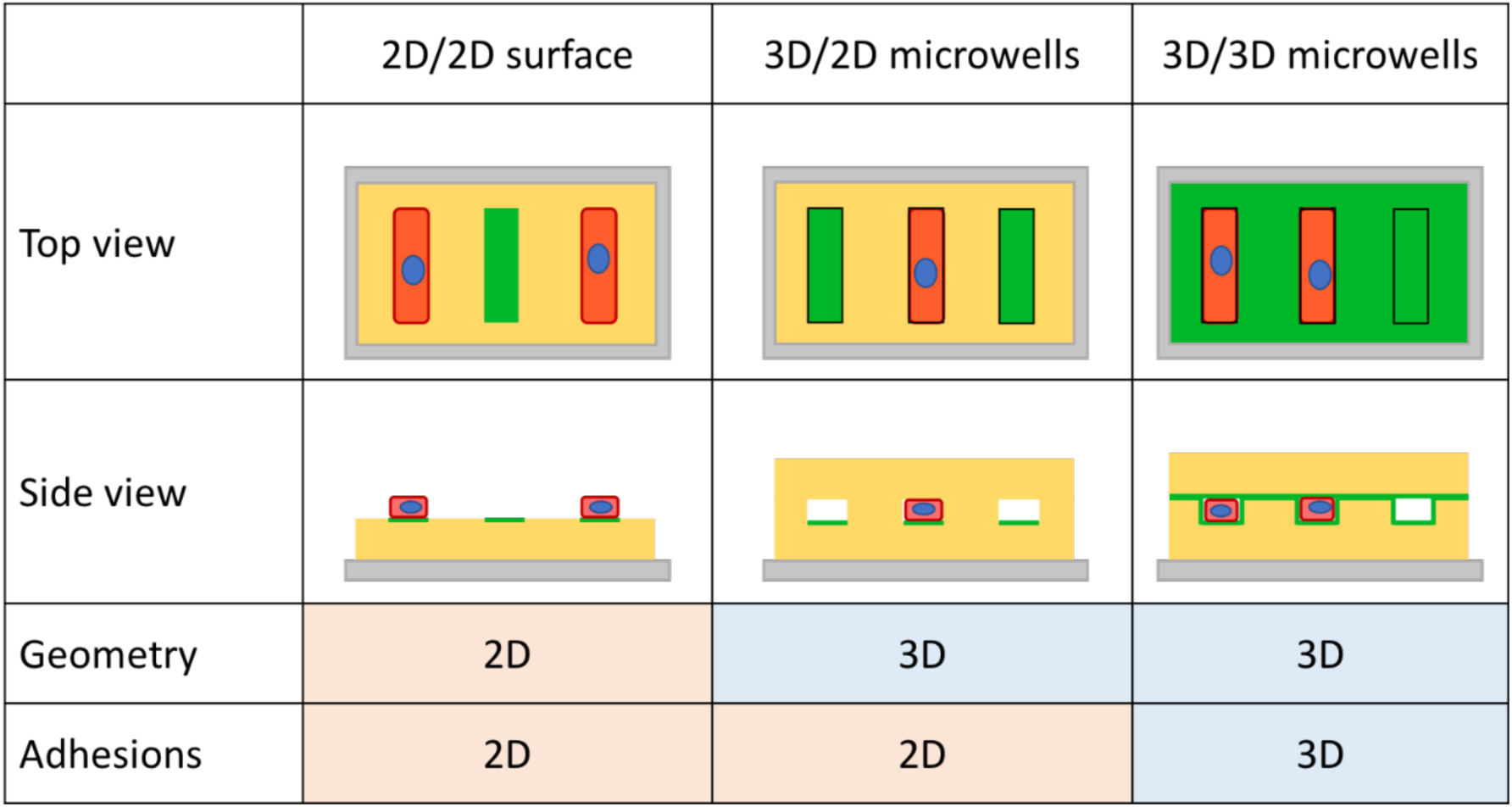
Schematic cross-section of three platform variations (gray: coverslip, yellow: polyacrylamide hydrogel, green: adhesive protein, pink: cell, blue: nucleus). The three platforms enabled decoupling of 3D confinement and the 3D adhesive environment, providing insight into what aspects of the 3D microenvironment most influence cell structure.

## Materials and Methods

### Fabrication of hydrogel platforms

We developed methods to fabricate three variations of single-cell polyacrylamide platforms: 2D/2D flat gels patterned with Matrigel rectangles, 3D/2D microwells selectively patterned with Matrigel to present the same 2D adhesive areas, and 3D/3D microwells fully coated with Matrigel (Figure 1). The overarching steps for fabrication were (1) making a Polydimethylsiloxane (PDMS; Sylgard 184) mold for the gel, (2) functionalizing the PDMS molds with protein to the desired locations, (3) casting the polyacrylamide gels on the PDMS molds, and (4) casting flat polyacrylamide gel lids (Figure 2).

**Figure 2:**
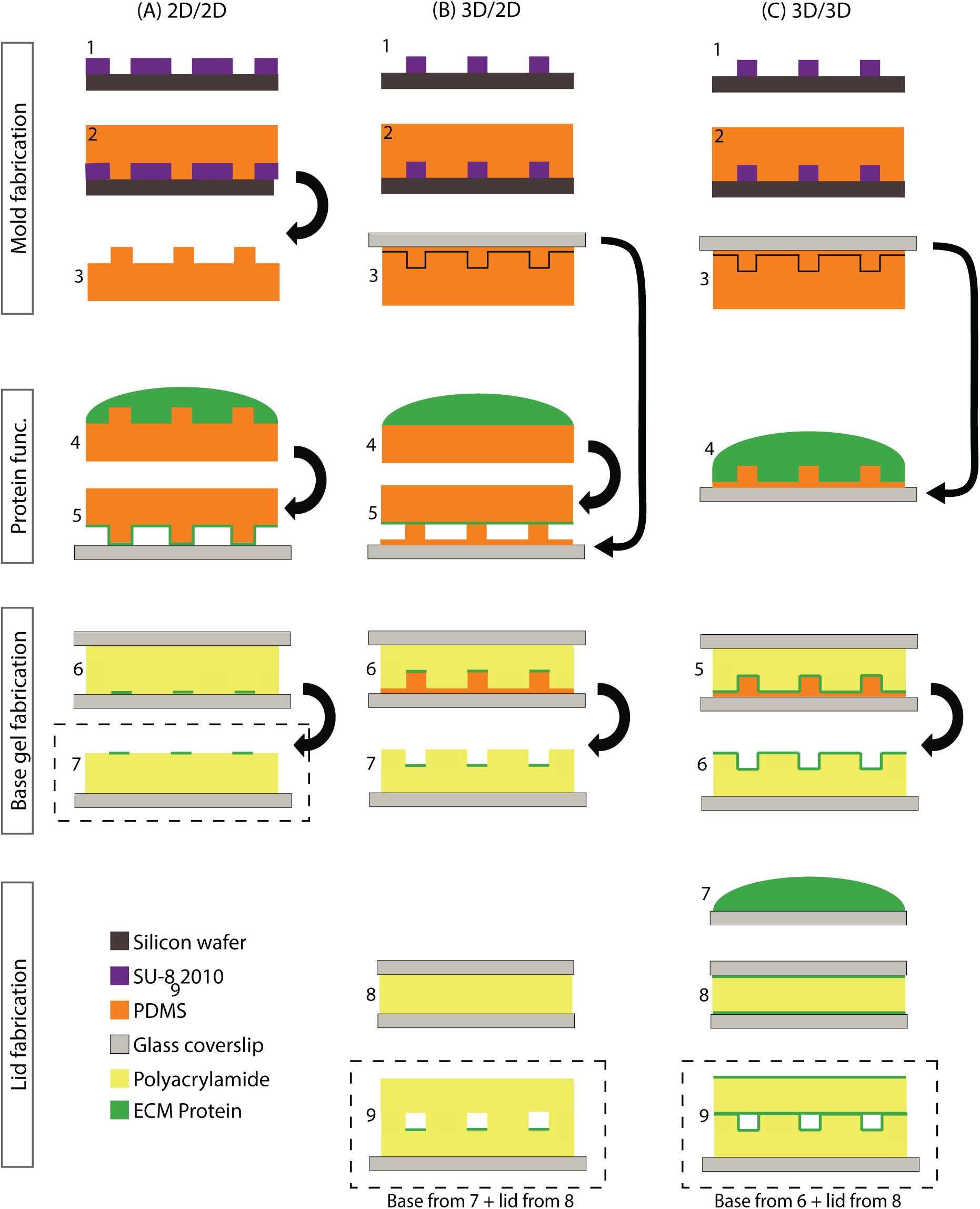
Microfabrication protocol for each polyacrylamide platform. **Mold/stamp fabrication:** we used SU-8 photolithography and PDMS molding to create 3D features used for microcontact printing and microwell molding. **Protein Functionalization:** we functionalized the polyacrylamide platforms with protein either through microcontact printing or by coating the mold surface. **Gel fabrication:** we cast polyacrylamide gels to create the three platforms, removing the top coverslip after polymerization. **Lid fabrication:** we created flat polyacrylamide lids, which we removed from the coverslips after polymerization and placed on top of the microwells to create a fully enclosed environment. The final platforms are outlined with dashed lines.

First, we fabricated SU-8 master molds to define features for both the 3D microwell molds and the 2D microcontact printing patterns (Figure 2: A1-C1). We used SU-8 2010 (Microchem) to achieve a feature depth of 12 μm using the protocol as written by the supplier and verified the height using profilometry. We used two transparency masks with the same pattern in opposite polarities, such that one could be used for the microwell molds and the other for 2D surface protein patterning. The features were rectangles with an aspect ratio of 5:1 and areas of 625 μm^2^, thus the microwell volumes were approximately 7,500 μm^3^. The features on the microwell wafer were raised from the surface, and the features on the protein patterning wafer were valleys. After the SU-8 was patterned, we silanized the wafers by vapor deposition with trimethylsilyl chloride (TMCS; Sigma 92361) for 1 hour to make a non-stick surface for molding.

For the 3D microwells, we made thin PDMS molds on coverslips by a double molding process (Figure 2: B, C 2-3). We poured, degassed, and cured bulk PDMS (Sylgard; Dow 4019862), mixed in a 5:1 ratio, to make the primary mold. We then diced each microwell mold and silanized them with TMCS for 1 hours. Next, we made a thin secondary PDMS mold for polyacrylamide casting. The thin PDMS mold was necessary to achieve repeatable geometries because thick PDMS has enough oxygen within the material to inhibit gel polymerization.^23^ To make the thin PDMS molds, we pipetted about 50 μL of PDMS (10:1 ratio) on the silanized primary PDMS mold and put an 18×18mm coverslip on top. We placed this mold assembly between two glass slides and applied pressure with a 50 g weight while curing.

We also poured PDMS (10:1 ratio) on the SU-8 master wafer for the 2D microcontact printing stamps (Figure 2: A2). After curing, we diced the PDMS into individual stamps (Figure 2: A3). We used regions with 3D PDMS features to functionalize 2D patterned gels and flat regions to functionalize 3D patterned microwells by microcontact printing.

The next step in the process was protein functionalization. We used gelatin, conjugated to Oregon green 488 (Invitrogen G13186) to visualize the protein patterning during development and characterization. We used growth factor reduced Matrigel (Corning 354230) for our studies with cells. We used Matrigel for its ability to support a broad-spectrum of cell-matrix interactions through its diversity of extracellular matrix components.^25^ In all cases, we incubated surfaces with Matrigel diluted in cold L15 media overnight at 4°C (10:1 dilution for microcontact printing 2D/2D and 3D/2D substrates and 30:1 for coating 3D/3D substrates). We used a higher concentration for microcontact printing than coating because protein transfer efficiency can be three times lower for microcontact printing.^36^ To functionalize coated microwells with protein, we simply incubated gelatin (100 μg/mL in PBS, 1 hour at room temperature) or Matrigel (1:10 dilution in L15 media, at least 6 hours at 4°C) on the thin PDMS microwell molds (Figure 2: C4). We also coated 18mm round coverslips with protein to use for casting the coated microwell lids (Figure 2: C7). For patterned substrates, we incubated gelatin (100 μg/mL in PBS, 1 hour at room temperature) or Matrigel (1:10 dilution in L15 media, at least 6 hours at 4°C) on PDMS stamps (Figure 2: A, B 4). We used flat PDMS stamps to functionalize 3D patterned microwell molds (Figure 2: B4) and molded PDMS stamps to functionalize 2D patterned gels (Figure 2: A4). Prior to casting the gels, we prepared the coated microwell molds by aspirating excess solution from molds. We also performed microcontact printing for both the 2D and 3D patterned substrates. To do this, we aspirated excess media from the stamps and dried them with nitrogen. We then plasma activated the substrates: flat coverslips for the 2D gels and PDMS microwell molds for 3D microwells. Next, we placed the stamps in contact with the appropriate substrates, topped each with a 50 g weight, and allowed to sit for 5 min (Figure 2: A, B 5).

To make the polyacrylamide gels, we first mixed and degassed polyacrylamide precursor solutions for 60 min. The precursor solutions contained acrylamide (Sigma 01696), bis-acrylamide (Sigma 146072), HEPES (Gibco 15630), and water. We added 2 μm red fluorescent FluoSpheres (Invitrogen F8826) for some studies during development to visualize 3D geometry of the polyacrylamide microwells. We characterized 12 polyacrylamide gel formulations with total polymer content (% T) of 5 – 12% and with crosslinker ratios (% C) of 1 – 3% (Table 1). For experiments with cells, we used the gel formulation for 8%T, 5 %C (10.6 kPa stiffness). We prepared base coverslips for the polyacrylamide gels using a bind silane protocol to allow the gel to bind to the coverslip.^21^ The final step was to cast the polyacrylamide gels. We initiated polymerization of each 500 μL gel precursor aliquot by adding 5 μL 10% w/v ammonium persulfate (APS; Sigma A9164) and 0.5 μL N,N,N”,N”-Tetramethylethylenediamine (TEMED; Sigma 411019). We cast each gel base with 45 μL and each lid with 35 μL solution. We allowed the gels to polymerize at room temperature in the dark for 40 min. After polymerization, we incubated the gels in PBS with 5% Penicillin-Streptomycin (Gibco, 15140122) overnight at 37°C.

**Table 1:** Polyacrylamide gel formulations used in this work, in microliters. Each formulation has a total volume of 1 mL but different total polymer contents (%T) and crosslinker ratios (%C).

| | Acrylamide<br>0.5 g/mL | Bis-acrylamide<br>0.025 g/mL | HEPES<br>250 mM | 2 μm<br>FluoSpheres | Water | APS<br>10%<br>(w/v) | TEMED |
| --- | --- | --- | --- | --- | --- | --- | --- |
| 7% T 1% C | 138 | 28 | 140.5 | 21.6 | 660.9 | 10 | 1 |
| 7% T 3% C | 136 | 84 | 140.5 | 21.6 | 606.9 | 10 | 1 |
| 7% T 5% C | 133 | 140 | 140.5 | 21.6 | 553.9 | 10 | 1 |
| 8% T 1% C | 159 | 32 | 140.5 | 21.6 | 635.9 | 10 | 1 |
| 8% T 3% C | 155 | 96 | 140.5 | 21.6 | 575.9 | 10 | 1 |
| 8% T 5% C | 152 | 161 | 140.5 | 21.6 | 513.9 | 10 | 1 |
| 10% T 1% C | 200 | 40 | 140.5 | 21.6 | 586.9 | 10 | 1 |
| 10% T 3% C | 194 | 122 | 140.5 | 21.6 | 510.9 | 10 | 1 |
| 10% T 5% C | 190 | 200 | 140.5 | 21.6 | 436.9 | 10 | 1 |
| 12% T 1% C | 238 | 48 | 140.5 | 21.6 | 540.9 | 10 | 1 |
| 12% T 3% C | 233 | 143 | 140.5 | 21.6 | 450.9 | 10 | 1 |
| 12% T 5% C | 1228 | 240 | 140.5 | 21.6 | 358.9 | 10 | 1 |

### Hydrogel swelling and stiffness characterization

We characterized geometric swelling and stiffness of molded polyacrylamide gels. We implemented custom image analysis (ImageJ) to measure microwell widths in brightfield micrographs. We used the relative width of the polyacrylamide microwell compared to design as a metric to quantify swelling of the geometry. To measure stiffness, we used atomic force microscopy (AFM) indentation. We used a 50 μm bead (Duke Scientific, 9050) attached to a cantilever (8.5 N/m stiffness; NanoWorld AG: Nanosensors, PPP-NCSTAuD-10) to indent the flat polyacrylamide surface between microwells by 0.6 μm, using methods developed previously in our group.^13^ We analyzed the force-distance curves using the Hertz elastic contact model to determine the gel’s elastic modulus, under the assumption that polyacrylamide is linearly elastic.^8,51^ We used R to plot mean values with error bars representing standard error of the mean.

### C2C12 culture, imaging, and analysis

We cultured C2C12 mouse muscle myoblasts (ATCC CRL-1772) in high glucose DMEM (ATCC 30-2002) and 10% FBS (Gibco 16000044) with 1% Penicillin-Streptomycin (Gibco, 15140122), and passaged the cultures until the confluency reached 70%. We chose myoblasts because muscle cells reside in 3D tissue and have a cylindrical or brick-like morphology *in vivo*.^7^ Thus, we expected a muscle myoblast to respond to a brick-like 3D environment, more so than cells that typically reside in monolayers and exhibit flattened morphologies such as endothelial cells. We seeded 30,000 cells per device, centrifuging at 300x-G for 1 min to increase the number of cells in wells, though for consistency, we centrifuged cells on all devices. We selected this seeding density as it was low enough to limit occurrences of multiple cells in microwells and high enough to provide multiple instances of single cells in microwells. Although we did not specifically optimize seeding efficiency, other work indicates that it may be possible to achieve at least 30% single cell efficiency, depending on microwell dimensions and cell type.^24,43^ We allowed the cells to recover at 37°C for about 20 min before adding the polyacrylamide lids to the microwell platforms, creating fully enclosed 3D environments.

Securing the PA lids in a fixed location relative to underlying microwell proved to be a key challenge. To keep the lids from moving, we created a custom laser-cut acrylic holder with magnets (Figure 3). This consisted of a 22×22mm acrylic sheet with 1.2 mm holes, spaced by 2 mm on a hexagonal lattice. We used acrylic (polymethyl methacrylate) because it has a high stiffness, providing enough structure to hold the polyacrylamide lid in place, and it is biocompatible.^19^ We glued 3×1 mm magnets (Amazon B07FYCRGBD) in the holder using PDMS, ensuring to coat the magnets entirely to prevent corrosion. We used the magnetic lid holder to secure the polyacrylamide lid by placing it on top of the microwell assembly and placing corresponding magnets underneath the cell culture plate (Figure 3). We allowed the C2C12 cells to adhere for 12 hours in the microwells using this setup prior to fixation in 4% paraformaldehyde (Fisher Scientific 50-980-487) for 20 min.

**Figure 3:**
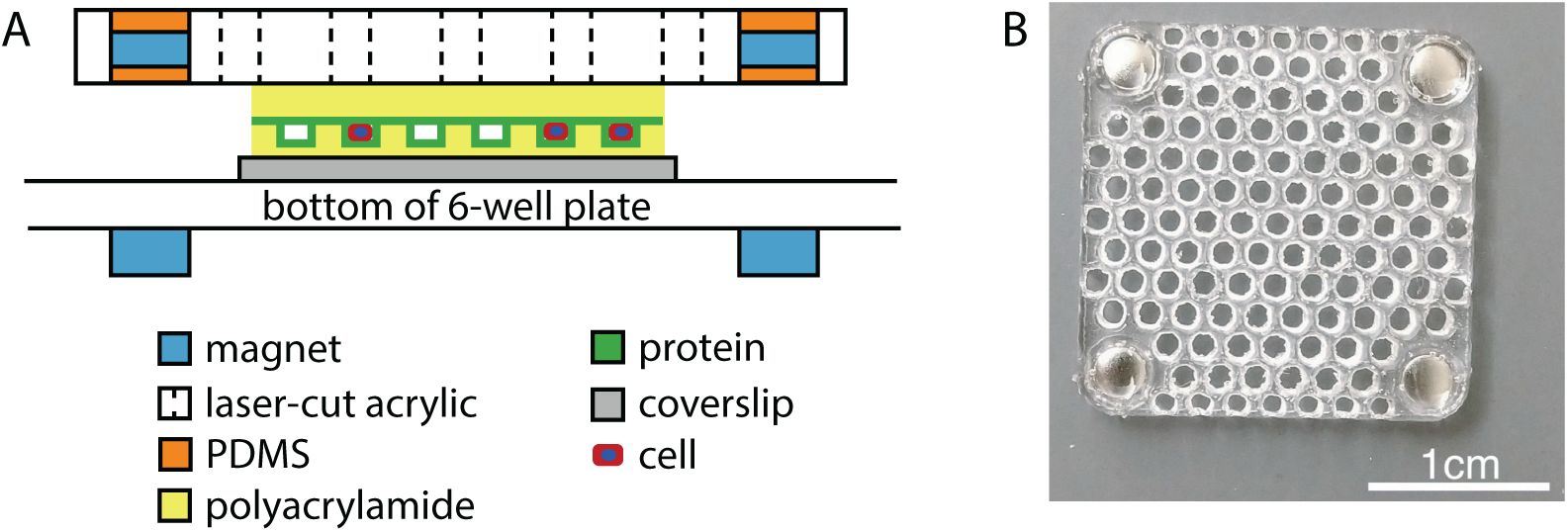
Laser-cut magnetic holder creates a stable, fully enclosed 3D environment for single cell culture. (A) Cross-section schematic of entire gel and lid assembly in a 6-well plate (not to scale). (B) Top view photograph of a laser-cut magnetic holder.

After fixation, we permeabilized the cells in 0.01% TritonX (Sigma X100-100ML) for 15 min. Next, we stained with ActinGreen 488 ReadyProbes (Invitrogen R37110) and 4′,6-diamidino-2-phenylindole (DAPI; Invitrogen D1306) for 30 min at room temperature.

We imaged the C2C12 cells on a Leica SP8 confocal microscope. Using FIJI,^46^ we performed custom analysis on the actin structures and nuclear shape per z-slice for each cell. To determine actin and nuclear height, we set a threshold for each stack individually and calculated the heights of the sub-stacks containing positive pixels at a value of at least 10% of the maximum slice. For the z-slice actin analysis, we measured the fluorescent intensity at each slice without processing. We calculated the orientation order parameter (OOP) for F-actin at each slice, using the equation

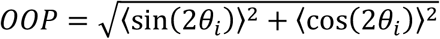

where *θ* is the fiber orientation at each pixel, *i*.^31^ Using this definition, OOP is 1 for perfectly aligned features and 0 for completely misaligned features. In practice, we applied the *Directionality* function in ImageJ to each slice after contrast enhancement. This function outputs the number of pixels with each orientation (−90° to 90° in 2° bins), which we input into the equation above to obtain actin OOP per z-slice.

To obtain 3D shape parameters for the nucleus, we implemented the ImageJ tool, 3*D Object Counter* on the binarized confocal stacks of the nuclei. This tool quantified the nuclear volume and sphericity. Sphericity, *ψ*, was calculated as

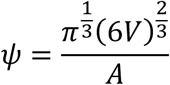

where *V* is the nuclear volume and *A* is the nuclear surface area. To determine the aspect ratio, we took the ratio of the major and minor axes after performing a z-projection.

We performed all statistical analysis and plotting in R. For all statistics, we used ANOVA to test for significant differences among our sample groups. If the ANOVA indicated significant differences, we used the Tukey HSD test to determine p-values between groups.

## Results

### Polyacrylamide formulation dictates swelling and elastic modulus

Prior to experiments, we defined and validated criteria for the hydrogel properties for cell culture. We wanted less than 10% swelling of the 3D geometry and a stiffness of approximately 10 kPa. Limiting swelling would allow us to fabricate more robust and repeatable microwell geometries, while a 10 kPa stiffness is physiologically relevant for muscle cells.^14^

We fabricated gels across a range of formulations and characterized them for swelling via normalized microwell width, and stiffness via AFM (Table 1). Normalized microwell width, which we used as a swelling metric, was highly influenced by crosslinker ratio and less by total polymer content (Figure 4 A-B). With a low crosslinker ratio of 1% C, the swelling was so great that the microwells were swollen shut regardless of total polymer content. With a high crosslinker ratio of 5%, swelling was minimal and the resulting microwell width was close to the design width, regardless of total polymer content. The range of normalized widths was 83 – 104% of design targets for all of our 5% C formulations. Thus, a crosslinker ratio of at least 5% C was most suitable for our application requiring achieving a targeted 3D geometry.

**Figure 4:**
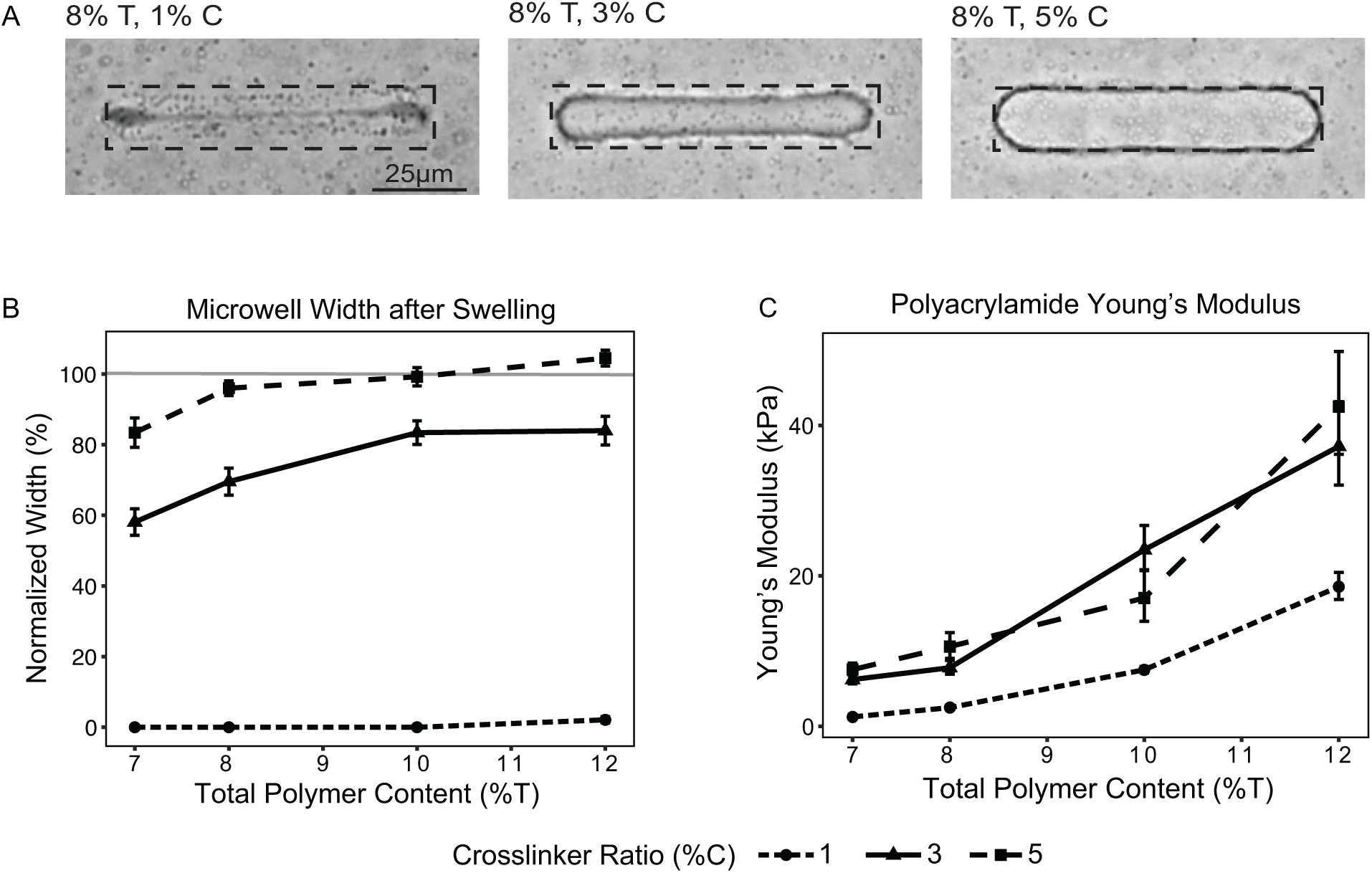
Polyacrylamide swelling and stiffness depend on the gel formulation. (A) Brightfield images of polyacrylamide microwells made with three different crosslinker ratios but the same total polymer content. Dashed boxes indicate the designed microwell geometry. (B) Geometric distortion of the 3D microwells induced by swelling, measured by normalized microwell width. A value of 0% indicates a microwell that is swollen shut, and a value of 100% indicates a microwell that has the exact dimension as the design (n=6 gels per condition, with each gel measurement taken as the average of 40 microwell widths; error bars show standard error). (C) Young’s modulus was measured using AFM. Measurements were made on the gel surface, in between microwells (n=5-6 gels per condition, with each gel measurement taken as the average of 25 stiffness measurements on each gel; error bars show standard error).

Next, we characterized the polyacrylamide stiffness using AFM indentation (Figure 4C). We found that both total polymer content and crosslinker ratio affected gel stiffness. Over our entire test range, we produced gels with Young’s moduli ranging from 1.4 to 47.7 kPa. Total polymer content was approximately linearly related to Young’s modulus over the range of formulations tested here. Increasing crosslinker ratio caused an increase in Young’s modulus as well, though this trend was strongest for gel formulations using ratios of less than 5% C. Comparing the Young’s modulus for gels with 3% C to 5% C for matching total polymer content showed no differences. For our experiments with cells, we chose to use an 8% T 5% C gel formulation because it exhibited our desired properties of limited swelling and a Young’s modulus of 10.6 ± 1.6 kPa.

### Fabricated platforms and C2C12 culture

Using the techniques described above, we successfully fabricated our desired microwell geometry and protein patterns for the three platform variations: 3D/3D, 3D/2D, and 2D/2D (Figure 5). Using a polyacrylamide formulation with 8% total polymer and 5% crosslinker ratio, we obtained platforms with an elastic modulus of 10.6 ± 1.6 kPa and with 3D geometries that were 96 ± 2% of the design geometries (measurements given as mean ± standard error).

**Figure 5:**
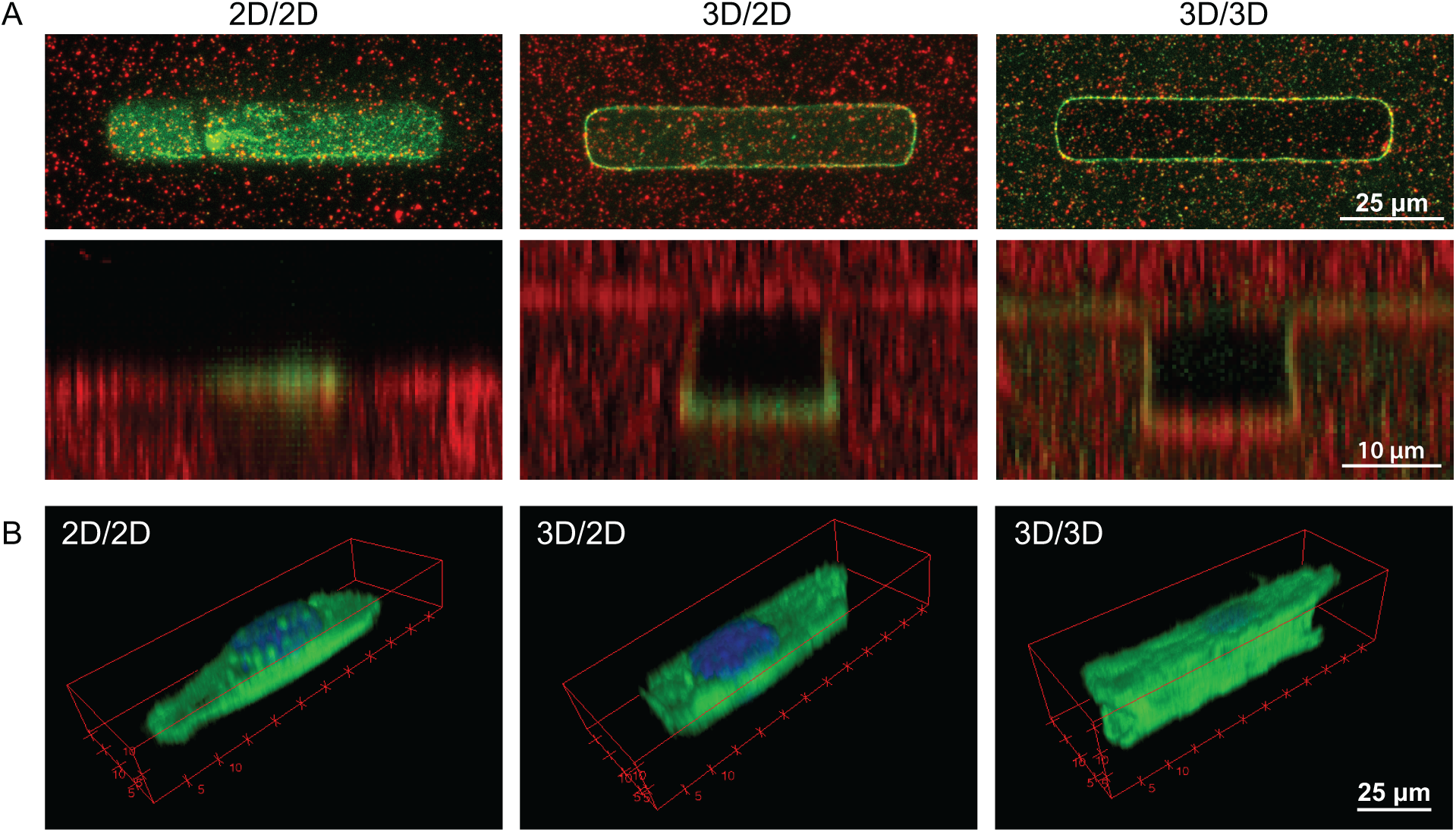
(A) Fluorescent confocal images of the three platform variations seen from the top view (top) and cross-sectional view (bottom) (green: fluorescent gelatin, red: embedded beads). Embedded beads were used for 3D visualization of the microwell geometry and fluorescent protein was used to confirm desired protein functionalization. (B) 3D renderings of single C2C12 cells on each platform. Each rendering was made from confocal slices of the cell (green: actin, blue: DAPI; wireframe box indicates image volume, not microwell volume).

We then cultured C2C12 cells on each platform for 12 hours. Initial experiments revealed that a polyacrylamide lid was critical to create an enclosed 3D microenvironment and to physically trap single cells within the coated microwells. Without the lids, the cells spread and migrated randomly on the 3D coated microwell platform instead of remaining in the wells, since the cells could attach to the entire platform surface. Additionally, the laser-cut magnetic holder was required to fix the polyacrylamide lid in place while still allowing for diffusion of oxygen and nutrients to the cells. Together, the polyacrylamide lid and laser-cut holder enabled us to create fully encapsulated 3D microenvironments and to trap single cells within wells. These technical advances allowed us to culture single C2C12 cells on 2D patterned (2D/2D), 3D patterned (3D/2D), and 3D coated substrates (3D/3D) (Figure 5).

### Effect of 2D and 3D geometry and adhesions on actin structure and nuclear shape

Inspection of the confocal stacks and 3D renderings indicated differences in the morphology and structure of the cells on different platforms. The cells on 2D/2D patterned gels appeared flat with the nuclei protruding from an actin base. Conversely, the cells in 3D/3D coated microwells had taller actin structures that filled the microwells and nuclei that were enclosed by the actin cytoskeleton (Figure 5).

To quantify these results, we analyzed F-actin intensity, F-actin OOP, and nuclear shape in single C2C12 cells across z-slices (Figure 6 and Figure 7). Using F-actin intensity data, we quantified the total height of F-actin in each cell. We found that cells confined on 2D/2D patterned gels were the shortest (median = 8 μm), followed by cells confined in 3D/2D patterned microwells (median = 9 μm), while cells in 3D/3D fully adhesive coated microwells had the tallest F-actin structures (median = 11 μm) and were statistically taller than cells on 2D/2D gels (Figure 6). We also examined the normalized F-actin intensity of the raw images (not binarized, split into three bins based on z-position). In this quantification of F-actin intensity, we found that intensity was consistent for cells on all three platforms from 0 to 9 μm in z-position. However, at the top third of the measured volume (10-15 μm), cells in the 3D coated microwells had over 2.5x F-actin intensity compared to both the 2D and 3D patterned platforms. These data suggest that while 3D confinement alone slightly increases the height of F-actin structures in C2C12 cells, an accompanying 3D adhesive environment supports much taller actin structures and a more 3D cell morphology.

**Figure 6:**
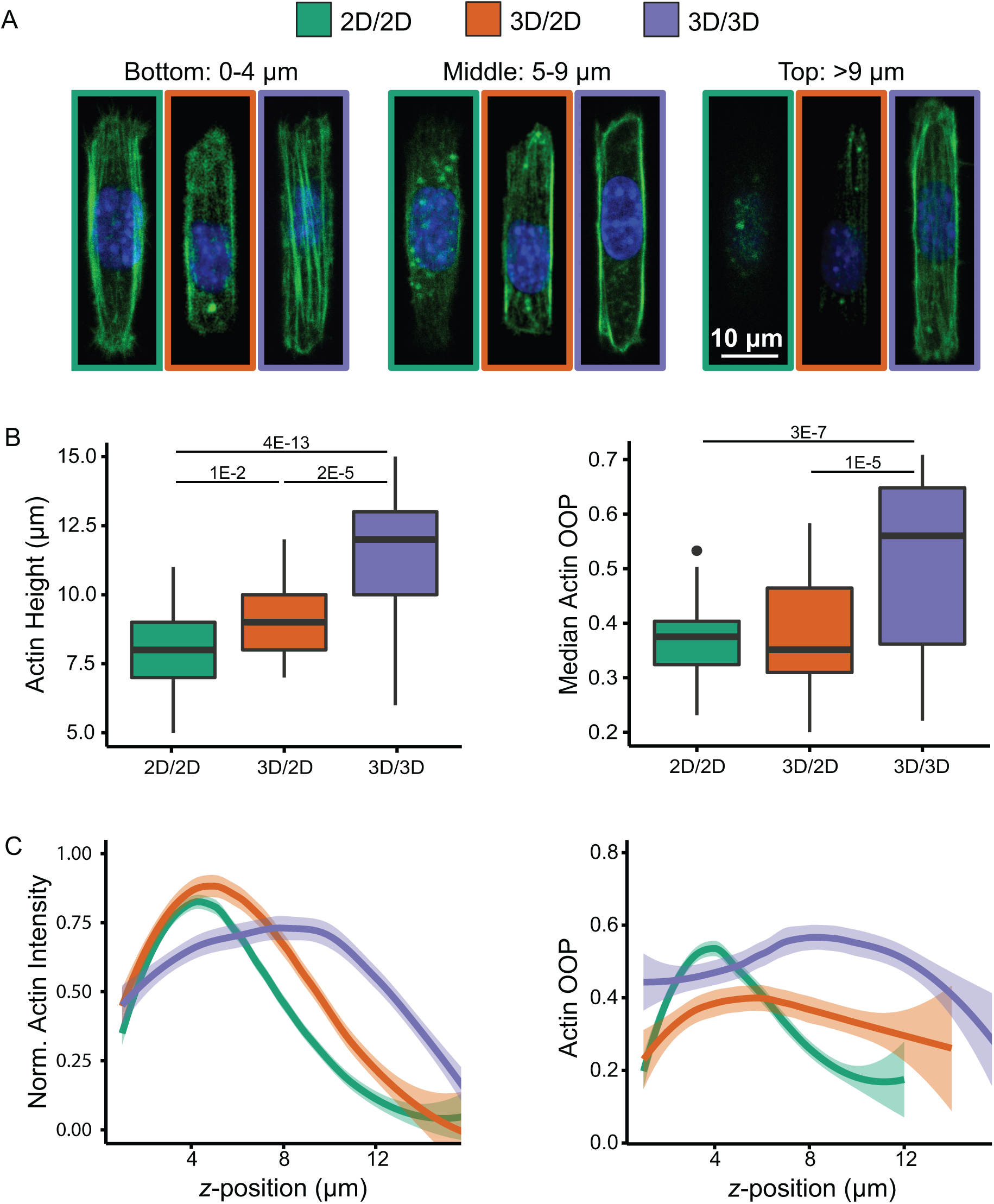
Slicewise actin analysis. (A) Representative confocal slices of cells on each platform at the bottom, middle, and top positions (green: actin, blue: DAPI). (B) Boxplots of actin height and median OOP in each cell. Boxplots show the minimum and maximum as whiskers, the interquartile range as the box, the median as the middle line, and outliers as points. P-values were calculated by ANOVA with post-hoc Tukey HSD Test (n > 28 cells per condition). (C) Normalized actin intensity and OOP for each confocal slice, fit with local regression curves and shaded with 95% confidence intervals.

**Figure 7:**
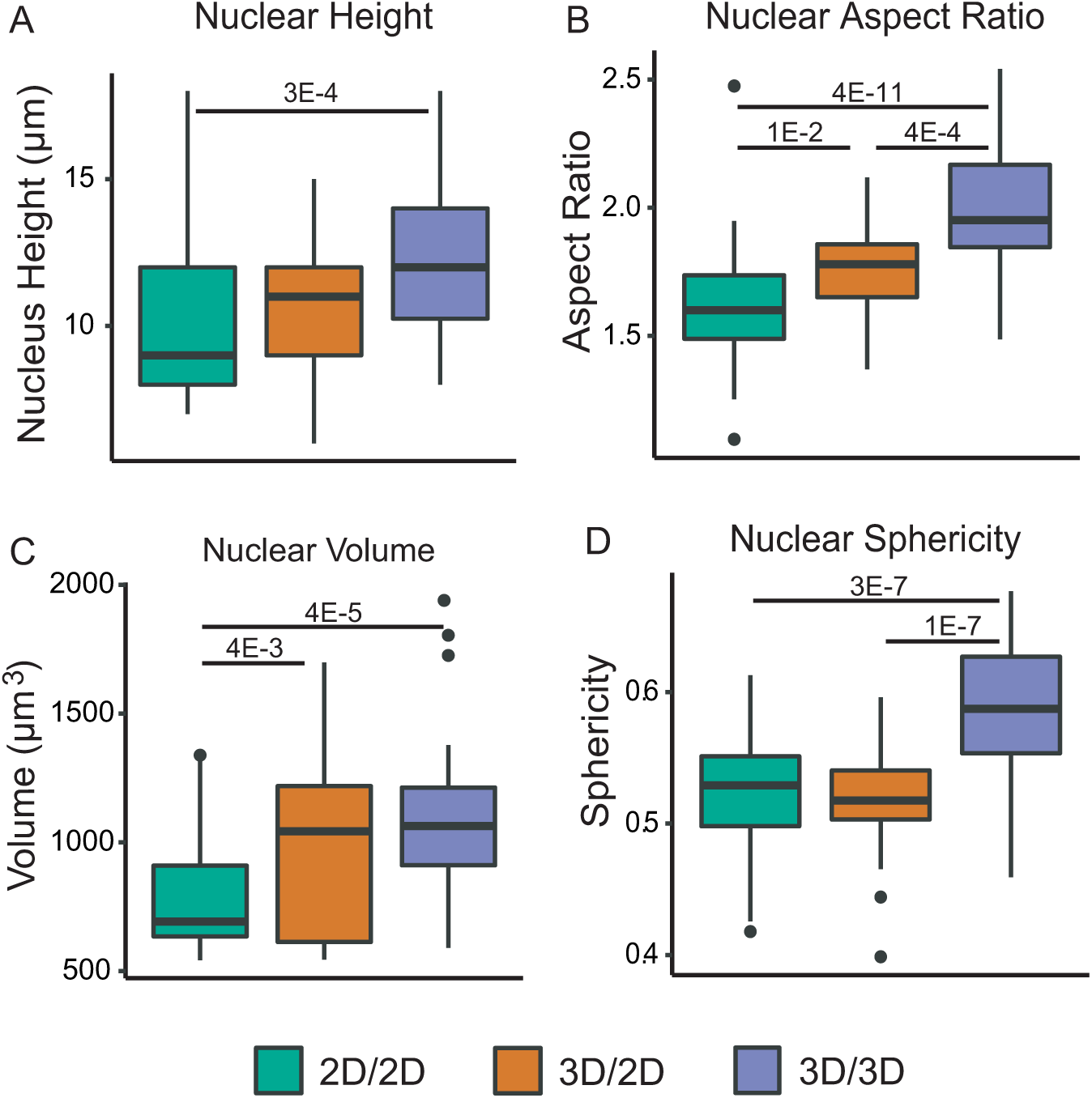
Nuclear shape analysis. (A) Boxplots of nuclear height. (B) Boxplots of nuclear aspect ratio, taken from a z-projection of each confocal stack. (C) Boxplots of nuclear volume. (D) Boxplots of nuclear sphericity. Boxplots show the minimum and maximum as whiskers, the interquartile range as the box, the mdian as the middle line, and outliers as dots. P-values were calculated by ANOVA with post-hoc Tukey HSD Test (n > 28 cells for all conditions).

Our analysis of F-actin OOP showed that 3D confinement and 3D adhesive environment, together, strongly increased F-actin alignment in C2C12 cells (p = 3E-7), while 3D confinement alone had no effect compared to the 2D control (Figure 6).Looking at the median OOP for each cell shows increased alignment in cells in 3D/3D coated microwells with a median OOP of 0.56. Comparatively, the median OOP for cells on 2D/2D patterned gels was 0.37, and the median OOP for cells in 3D/2D patterned microwells was 0.35. Interestingly, when we examined the data across *z*-slices, we found that the F-actin organization with respect to *z*-position were quite different for the three platforms. Cells in both the 3D/3D coated and 3D/2D patterned microwells had fairly consistent actin OOP through the *z*-position. In contrast, cells on 2D/2D patterned gels had a peak in F-actin OOP at a *z*-position of 4 μm, but then showed a rapid decrease in organization with increasing *z*-position. These data suggest that 3D confinement is sufficient to drive a more uniform F-actin organization in *z*. However, cells exposed to 3D confinement alone (3D/2D patterned microwells) still had lower OOP overall than cells exposed to 3D confinement and 3D adhesive environment (3D/3D coated microwells).

We also found that 3D microenvironments altered nuclear geometry (Figure 7). 3D confinement alone (3D/2D patterned microwells) resulted in a significantly increased nuclear aspect ratio along the microwell’s long axis and volume compared to cells on 2D/2D platforms. Cells in 3D/3D coated microwells also showed increased nuclear aspect ratio and volume, as well as increased height and sphericity. The median nuclear height was 9 μm for cells on 2D/2D patterned gels, 11 μm for cells in 3D/2D patterned microwells, and 12 μm for cells in 3D/3D coated microwells. Similarly, the median nuclear aspect ratio increased from 1.60 for cells on 2D/2D patterned gels, 1.78 for cells in 3D/2D patterned microwells, and 1.95 for cells in 3D/3D coated microwells. Thus, 3D confinement tends to elongate and heighten nuclei, which is reflected by the increased volume and sphericity. These results followed the same trends observed in actin structure, showing that 3D confinement alone influences nuclear structure, and the addition of a 3D adhesive environment drives more dramatic changes in nuclear shape.

## Discussion

To decouple the effects of 3D confinement from 3D adhesive environments, we developed and deployed novel polyacrylamide microwell platforms. One challenge in platform development was minimizing polyacrylamide swelling to achieve our desired microwell geometries. Bulk polyacrylamide swelling has been studied extensively, as a measurement of the mass of water in an equilibrated gel.^39^ However, relatively little research has explored the effects of swelling on cast 3D polyacrylamide features. Our results agree with previously published results, showing that geometric swelling decreases with crosslinker ratio but is not highly affected by total polymer content.^11^ The effects of gel formulation on elastic modulus from the literature are somewhat unclear as they can be highly affected by measurement technique.^13,39^ However, metadata analyses indicate that total polymer content increases modulus while crosslinker ratio may also increase modulus modestly, which agrees with our results.^13,39^ Thus, by maintaining a high crosslinker ratio and varying total polymer content, we could limit swelling while tuning elastic modulus to match our biological application.

Compared to other similar single cell microwell platforms, the most important features of our system are the material properties and fully enclosed 3D microenvironment. Other single cell microwell environments have been fabricated with PDMS,^38,50^ methacrylated hyaluronic acid gels,^4^ or polyethylene glycol gels.^50^ These materials have tunable mechanical properties but are not linearly elastic. Our microwells are molded from polyacrylamide, which behaves as a linearly elastic material under our experimental conditions.^8,42^ Thus, in the future, it would be possible to extract cellular forces by measuring displacements of embedded fluorescent beads within the microenvironment and using, for example, finite element methods.^18,28^ Such analyses would provide interesting insights into how changes in the 3D microenvironment could influence cellular contractility.

Another important, novel feature in our microwell platform is the polyacrylamide lid, which maintains a stable, fully 3D environment and can be coated with protein if desired. We found that securing the polyacrylamide lid with the magnetic holder was essential for maintaining a stable microenvironment, while allowing for diffusion of gases and small molecules. Other studies had success with an unsecured lid setup, consisting of a thin hydrogel layer sitting on top of the microwells.^3,4^ Although we tried this technique, we found it difficult to ensure the devices were submerged in adequate media without the lid moving around. A novel, magnetically attached lid holder placed on top of a free polyacrylamide lid allowed us to create a stable, fully enclosed 3D microenvironment for single cells.

Studying C2C12 cells on the polyacrylamide platforms gave us insight into how 3D confinement and 3D adhesive environment each affect intracellular structure. A number of studies have shown that 2D cell adhesive patterning of cell shape and spread area can affect cytoskeletal structure.^9,52^ Our findings indicate that 3D confinement alone is sufficient to affect both actin and nuclear organization. The addition of a 3D adhesive environment increased the magnitude of these structural effects. Recent microtissue engineering studies have also found that cells in 3D environments are heterogeneous in size, have varied morphologies, and exhibit different morphologies and gene expression profiles compared to cells in 2D monolayers.^37,45,49,54^ The platforms developed here will thus enable more controlled studies of the effects of 2D and 3D confinement and mechanosignaling in cells to provide new insights for microtissue designs to achieve desired cellular geometries.

An open area of investigation is how changes in nuclear shape affect chromatin arrangement and gene expression. Previous work has shown that the cytoskeleton can interact with the nucleus changing its shape, volume, chromosome arrangement, and may influence gene expression.^12,27,33,41,56^ Actin and microtubule interactions with the nucleus have been the focus of many studies investigating nuclear deformation.^27,30,41,56^ Interestingly, studies utilizing 2D micropatterns have found that highly confined cells had shorter actin filaments, softer nuclei, and more dynamic chromatin, suggesting that the cytoskeletal network plays a role in nuclear structure and gene expression.^33^ Recent work studying breast cancer cell migration through constricted microchannels also measured extremely high nuclear deformability, mediated by interactions between actin and the nuclear envelope via lamins and Arp2/3 actin nucleation surrounding the nucleus.^20,22,53^ The nuclear deformation we observed in our 3D platforms could be influenced by these interactions and this would be an interesting area of future work. Such interactions could be probed within our 3D platform using pharmacological agents to interfere with actin polymerization. Such studies could provide new insight into how the nucleus and cytoskeleton interact within a 3D microenvironment. In addition, it would be interesting to investigate if our novel 3D environment could cause changes in chromatin arrangement and gene expression, since our platform allows for control of geometry, protein adhesion, and even substrate stiffness.

Delving further into single cell analysis, for instance single cell gene expression could help determine mechanisms for the responses in intracellular structure demonstrated in this work. Downstream assays like RNAseq would require significant modification to the platform in order to enable harvesting of single cells for pairwise RNAseq. However, alternative approaches like fluorescence in situ hybridization (FISH) may be possible in the platform described here with fewer modifications.

Other interesting areas for future work could include studying different cell types in these platforms or utilizing different proteins for surface functionalization. Investigating how different cell types respond to our 3D microenvironments would reveal which structural changes represent conserved biophysical responses, which depend on directional contractility characteristic of a subtype of cells like muscle cells, or which depend on specific integrin subtypes. Similar studies indicate that fibroblasts^38^ and mesenchymal stem cells^3,4^ alter their cytoskeleton and nuclei in response to single-cell 3D microenvironments; however more work is needed to quantify the responses in 3-dimensions and investigate other cell types. Utilizing different proteins for surface functionalization would also be a fascinating area of future research. From 2D studies, it is clear that both ECM composition and ligand density can regulate numerous cellular processes through outside-in signalling.^34,47,48^ For instance, cytoskeletal proteins can interact with integrins through intermediate proteins, altering intracellular structure.^10^ In addition, signaling molecules can be sequestered in adhesion complexes, affecting gene expression. Future work is needed to fully characterize whether ligand type and density are differentially sensed in 3D vs. 2D configurations.

Here, we found that C2C12 cells in 3D/3D coated microwells exhibited F-actin organization that differs from cells on 2D/2D patterned gels, and thus we would expect differences in actin-nuclear interactions. For instance, in 2D, an actin perinuclear cap is thought to largely impact nuclear mechanics.^27,30,41^ However, in our 3D/3D coated microwells, the F-actin organization (intensity and OOP) was consistent along the z-axis and we did not observe the actin take on a perinuclear cap structure. The F-actin structures observed in 3D/3D environments likely interact the nucleus differently than the actin cap seen in cells on 2D/3D patterned gels. Our platform will enable future studies to examine how a 3D microenvironment affects the structure and interplay of such intracellular signaling.

## Conclusions

Here we presented a method to make polyacrylamide platforms to systematically manipulate cell morphology in tunable 2D and 3D microenvironments. By altering the gel formulation, we found that we could control the geometric swelling and elastic modulus of polyacrylamide hydrogels independently. In addition, using protein micropatterning techniques, we were able to functionalize both 2D surfaces and 3D microwell platforms using microcontact printing. This allowed us to create three platform variations, effectively decoupling the effects of 3D confinement and 3D adhesive environment. C2C12 cells grown on these platforms demonstrated that 3D confinement alone altered F-actin orientation and nuclear shape. The combination of 3D confinement and a 3D adhesive environment resulted in more distinct changes in F-actin and nuclear structure including increased F-actin alignment and height and increased nuclear aspect ratio. This platform will enable single-cell studies investigating the effects of 3D microenvironments on intracellular structure and downstream mechanosignaling effects.

## Acknowledgements

This work was supported in part under grants NIH RM1 GM131981, AHA 17CSA33590101, NIH UG3 TR002588. In addition, R.E.W and A.K.D would like to acknowledge graduate research funding through the Stanford Graduate Fellowship, Stanford DARE Fellowship, and NSF GRFP. A.R.D. gratefully acknowledges support from the NIGMS (1R35GM130332) and Howard Hughes Medical Institute (Faculty Scholar Award)

